# GeneDART: Extending gene coverage in image-based spatial transcriptomics by deep learning-based domain adaptation with barcode-based RNA-sequencing data

**DOI:** 10.1101/2023.02.07.527488

**Authors:** Jungyoon Ohn, Daeseung Lee, Hongyoon Choi

**Affiliations:** Portrai, Inc., Seoul, Republic of Korea; Department of Nuclear Medicine, Seoul National University Hospital, Seoul 03080, Republic of Korea; Department of Nuclear Medicine, Seoul National University College of Medicine, Seoul 03080, Republic of Korea

## Abstract

Spatial transcriptomics (ST) technologies provide comprehensive biological insights regarding cell-cell interactions and peri-cellular microenvironments. ST technologies are divided into two categories: imaging-based (I-B) and barcode-based (B-B). I-B ST technologies provide high resolution and sensitivity but have limited gene coverage. B-B ST technologies can analyze the whole transcriptome but have lower spatial resolution. To address these limitations, we propose a deep learning-based model that integrates I-B and B-B ST technologies to increase gene coverage while preserving high resolution. A model, trained by a neural network with an adversarial loss based on I-B and B-B datasets from human breast cancer tissue, was able to extend gene coverage to whole transcripts-level and accurately predict gene expression patterns in the I-B dataset with a high resolution. This novel methodology, named GeneDART, could enable researchers to utilize B-B and I-B ST datasets in a complementary way.

## Introduction

Spatial transcriptomic (ST) technologies provide gene expression landscapes of biologic tissues to understand cells in their local microenvironment and cell □ cell interaction context. (1,2) The technologies for ST could be divided broadly into two categories: imaging-based (I-B) and barcode-based (B-B) technologies. (3,4) I-B technologies encompass *in situ* hybridization and *in situ* sequencing methods, which are prominent for their high spatial resolution and sensitivity capable of detecting messenger RNA (mRNA) at subcellular level. (2) Considering that a specific mRNA associated transcription, trafficking and localization information within a cell provides an important clue for cell-cell interaction in its niche (5), their ability to localize mRNAs in subcellular resolution could enable scientists to deeply understand biological processes. However, some technical points, such as a long microscopic imaging time with generating massive data for relatively small number of targeted genes, limit the versatility of I-B technologies in ST. (6,7) Meanwhile, B-B technologies represented by array-methods are widely used to investigate the unbiased whole transcriptome by next-generation sequencing process for polyadenylated mRNA in larger tissue specimens. (3) Though, its technical dependency on a fixed array makes it difficult to track each transcript for the individual cellular level, which is resulting in lower spatial resolution. (2) This drawback restricts researchers from using B-B technologies to verify the biological features in a cellular level, one of the ultimate goals of ST. (3) In this respect, I-B and B-B technologies could be considered as complementary relationships with clear limitations of each. To overcome the limitations of each ST method, we herein propose a deep generative model of ST data by conjoining both technologies, by which we could extend gene coverage of I-B ST with a highly preserved spatial resolution.

## Materials and Methods

### Data collection and preprocessing

In this study, we used I-B ST dataset (https://www.10xgenomics.com/products/xenium-in-situ/preview-dataset-human-breast) and B-B ST dataset (https://www.10xgenomics.com/resources/datasets/human-breast-cancer-ductal-carcinoma-in-situ-invasive-carcinoma-ffpe-1-standard-1-3-0) from human breast cancer tissue generated by Xenium and Visium platform technology (10x Genomics, CA, USA), respectively. The expression matrix and metadata of cell type information including location in I-B dataset were loaded using scanpy (version 1.6.1). (8) As a preprocessing step, both data were normalized by dividing the counts within each cell by the total number of transcripts, scaled by 10^6^, and then log-transformed. Some of the genes commonly present in both dataset (I-B and B-B ST) were selected for the input (i.e. *Genes for Input*) of a deep learning-based model (GeneDART model) (Fig. 1A).

**Figure 1.**
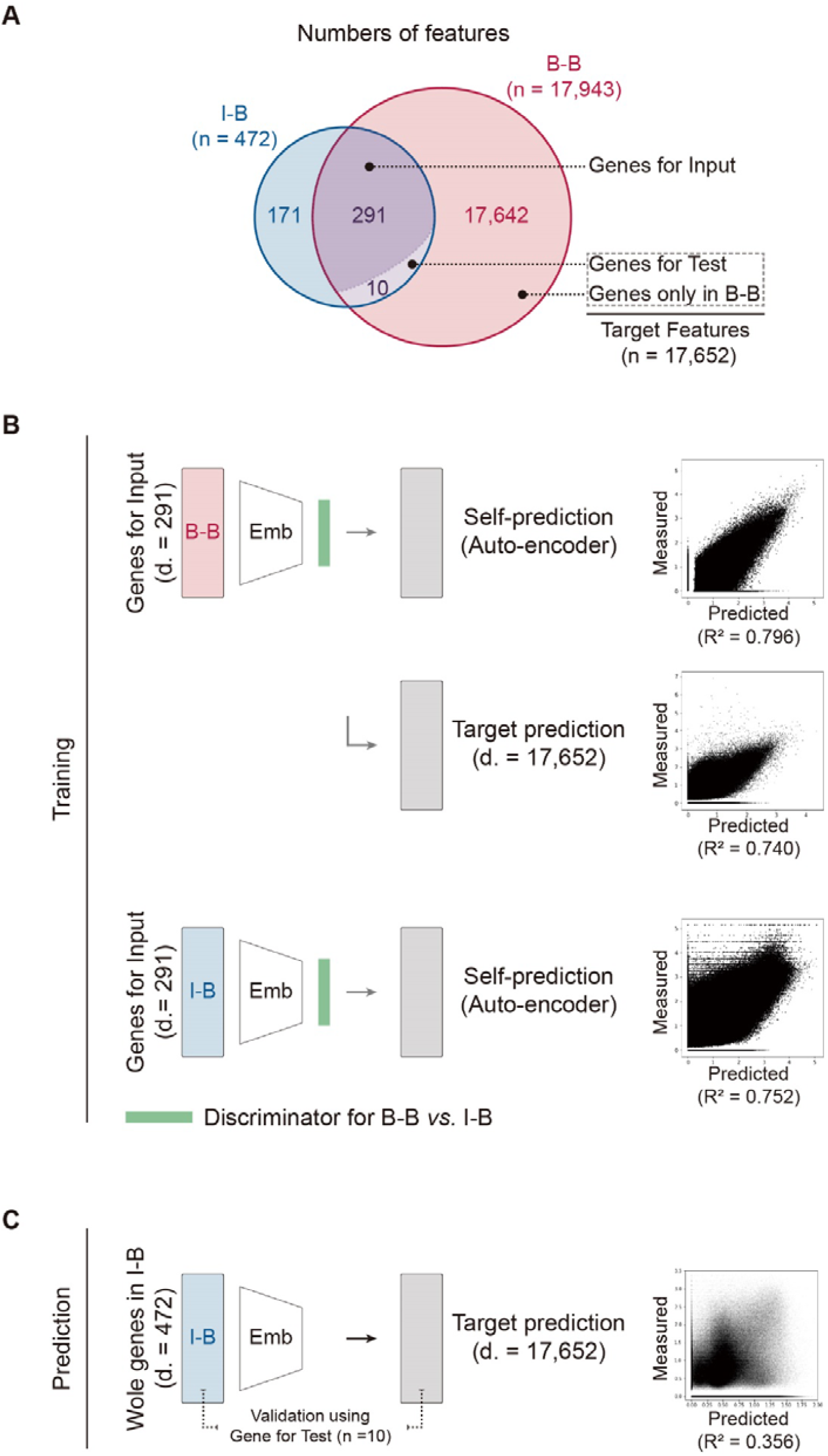
Extending gene coverage in I-B spatial transcriptomics by deep learning-based domain adaptation with B-B RNA-sequencing data. (A) Diagram for defining the gene sets: *Genes for Input, Genes for Test*, and Genes only in B-B. (B) Training of a neural network attached with an adversarial loss. (C) Extending gene coverage in I-B by the established neural network model, GeneDART. **Abbreviation**: B-B, barcode-based; d., dimension; Emb., Feature embedder; I-B, imaging-based; ST, Spatial transcriptomics.

### Training of GeneDART model

The GeneDART model consisted of a feature embedder and a discriminator, which were neural networks. The outputs were connected to the embedded features and resulted in ‘Self-prediction’ or ‘Target features-prediction’. The ‘self-prediction’ meant predicting the input features, which is similar with auto-encoder. Thus, in this model, it predicted expression data of ‘*Genes for Input*’. The ‘Target features-prediction’ was for the gene expression data of ‘Target features’: genes that randomly selected ten common genes (*Genes for Test*) in both the B-B and I-B ST datasets and included only in the B-B ST dataset (Fig. 1A). The feature embedder had two layers composing 128 dimensions. Batch normalization and ReLU activation were used for each layer. Another neural network, discriminator, was connected to the embedded features to discriminate on whether input was originated from the B-B or I-B ST dataset. The discriminator had another layer with 128 dimensions and then connected to one-dimensional output with sigmoid activation.

The loss function of the gene feature prediction was defined by mean squared error (MSE). The loss was calculated by two outputs, ‘self-prediction’ and ‘Target features-prediction’. In addition, to make adversarial loss, binary cross-entropy, a probability that a certain input was correctly allocated to the assigned input type, B-B versus I-B.

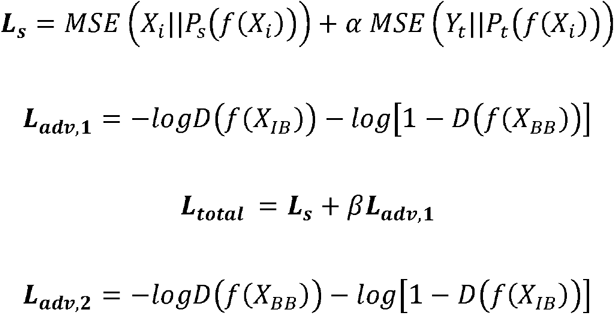

*X*_*i*_: *Input gene features, X*_*IB*_: *input I-B gene features, X*_*BB*_: *input B-B gene features, Y*_*t*_: *Target gene features, prediction for unknown genes, P*_*s*_: *gene expression prediction model for ‘Self-prediction’, Pt: gene expression prediction model for ‘Target features-prediction’, D: discriminator model for B-B vs. I-B, α: weight of loss between Self-prediction and Target features-prediction, β: weight of loss between MSE loss and adversarial loss*.

To begin the training process (Fig 1B), this model was initialized using an auto-encoder of the I-B data and the *Ls* loss with a value of alpha equal to zero. After this pre-training, the optimization was done using an adversarial loss, which involves training the model to minimize the standard loss function with inverted labels. Two optimization processes were then applied. The first process involved training the networks to minimize both *L*_*s*_ and *L*_*adv,1*_, while keeping the weights of the discriminator fixed. The second process involved training the discriminator to minimize *L*_*adv,2*_, while keeping the weights of the feature embedder, *f*, fixed. These two processes were repeated a certain number of times, as specified by a training parameter. The parameters including iteration number of training were determined by predicted output of the ten randomly selected common genes (*Genes for Test*). Then, the finally trained GeneDART model was applied on the I-B ST dataset to predict other genes not included in the dataset, extending gene coverage (Fig 1C).

### Validation of GeneDART model

To validate the GeneDART model, the ten randomly selected genes among the common genes (*Genes for Test*) were used, based on that these genes were not included in the *Genes for Input*, but included in both the predicted output by the GeneDART model and measured values in the I-B ST dataset. *Genes for Test* were inputted and then ‘Target features-prediction’ was performed to generate the predicted data (Fig. 1C). The output matrix was then incorporated into the scanpy object for further visualization and analysis. The predicted data and the measured data in the I-B ST dataset were compared to check whether this model predicts gene expression patterns reliably.

## Results

The imported I-B ST dataset consisted of a total of 167,782 cells in a single field-of-view with gene expression data of 472 features. To extend coverage of gene expression data up to the level of the B-B ST dataset which contains 17,943 gene features, we firstly sorted out the common genes in both B-B and I-B dataset (301 genes), by randomly categorizing them into two groups: 291 genes (i.e. *Genes for Input*) and 10 genes (i.e. *Genes for Test*) (Fig. 1A). The *Genes for Test* and the genes only in B-B dataset (17,642 genes) were designated as *Target features* (17,652 genes) (Fig. 1A). The aim of the model was to predict the *Target features* using *Genes for Input*. To solve the difference between B-B and I-B data in terms of domain, an adversarial loss was added for domain adaptation. Specifically, a neural network for discriminating I-B and B-B ST datasets was constructed to define adversarial loss. The models were optimized by tuning their parameters to minimize the sum of the loss in the prediction values for (1) the *Genes for Input* itself in the B-B dataset and (2) the *Target features* in the B-B dataset, and (3) the *Genes for Input* in the I-B dataset as well as adversarial loss (Fig. 1B). As a result, the trained neural network showed its performance as R^2^ = 0.796 for self-prediction (auto-encoder part that predicts *Genes for Input*) and R^2^ = 0.740 for Target features in the B-B dataset (Fig. 1B). In the I-B dataset, its self-prediction (auto-encoder) performance was gauzed as R^2^ = 0.752. The established neural network model was then used to the I-B ST dataset for extending the gene coverage in I-B ST dataset to the level of B-B ST dataset (Fig 1C).

To prove the accuracy of the proposed method, the predicted data and the measured data in the I-B ST dataset of *Genes for Test* (randomly selected 10 genes) were compared, resulting in statistics by Spearman’s correlation coefficient (*ρ*) ranging from 0.107 to 0.700 (*p <* 0.001) (Fig. 2A). The predicted values were also fitted with the B-B ST dataset (*ρ* ranging from 0.255 to 0.694, *p <* 0.001) (Fig. 2B). The gene expression patterns with spatial information of each gene in *Genes for Test*, which were re-portrayed based on the predicted dataset, were compared to those in the I-B ST dataset respectively (Fig. 2C). The predicted up- or down-regulated gene expression patterns on the tissue histology maps were compatible to the spatially distributed signals based on the I-B ST dataset.

**Figure 2.**
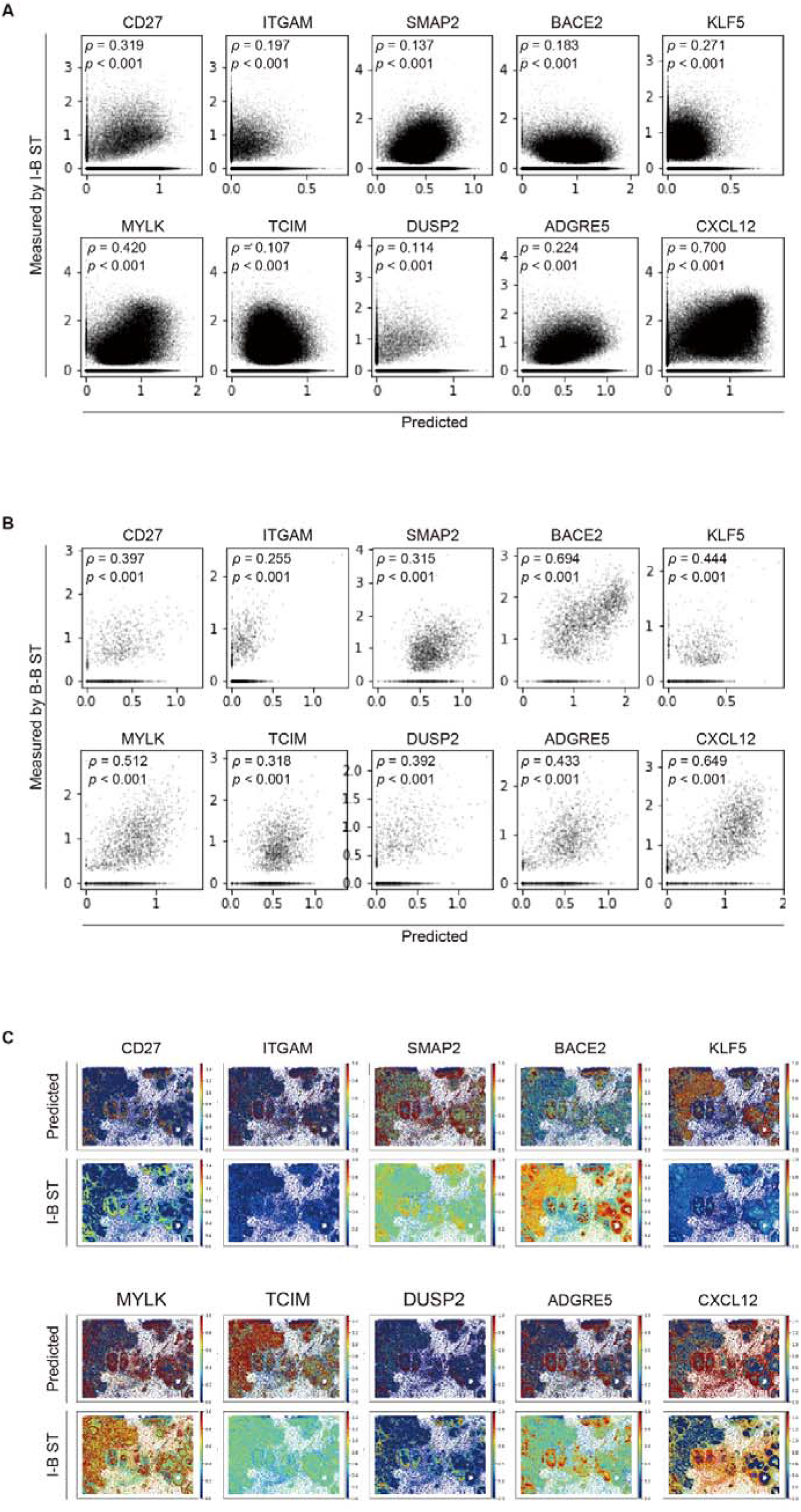
The accuracy and consistency of the GeneDART model. The statistics of the association between the measured values in I-B dataset (A) or in B-B dataset (B) and the predicted values by the neural network model of each randomly selected 10 genes (*Genes for Test*), based on the Spearman’s correlation coefficient (*ρ*). (C) The spatial gene expression patterns of *Genes for Test*, based on the predicted by the neural network model (upper panel), and the I-B ST dataset (lower panel), respectively. **Abbreviation**: B-B, barcode-based; I-B, imaging-based,

## Discussion

The spatially well-coordinated cells in organs, such as brain, kidney, liver, and gastro-intestine, and the appropriate gene expression in those cells are crucial for maintaining biological functions. Furthermore, an important disease such as cancer shows intratumoral spatial transcriptomic heterogeneity across the cells in the cancer tissue, and this heterogeneity is associated with the therapeutic response and prognosis of the disease. (4,9,10) Likewise, mapping the transcriptomics interface between cancer and its microenvironment enables researchers to understand the progression of diseases comprehensively. (11,12) In this respect, it is crucial to obtain information regarding the quantification of mRNA transcripts as proxies for a specific gene expression in the context of spatial information, in order to explore life science and investigate pathomechanisms of diseases. (2)

Numerous technologies for ST have been continuously developed and introduced for decades, as systematically described in recent articles. (2,3) However, considering all the aspects of the resolution of mRNA detection in tissues on slides and the number of genes that can be quantified in terms of cost and time efficiency, no specific ST technology is superior. Broadly, I-B and B-B technologies have advantages in the context of spatial imaging resolution and the number of coverage genes that can be identified, respectively. Accordingly, researchers choose appropriate techniques, depending on the purpose of the experiment. (2,3) In this respect, establishing a computational processing pipeline that can broadly disclose transcriptomes in local cells at a high resolution could enable researchers to be able to maximize their research efficiency.

As part of this effort, several researchers introduced a method for integrating single cell RNA-sequencing (scRNA-seq) dataset with ST dataset. Along with scRNA-seq, a high-resolution sequential fluorescence in situ hybridization (SeqFISH) method could demonstrate the spatially resolved expression of genes that are not detected by seqFISH in a cellular resolution. (13) Rodriques *et al*. integrated the cell type signature from scRNA-seq with Slide-seq data could facilitate the discovery of spatial gene expression patterns. (14) Furthermore, comprehensive analysis of multiplexed ion beam imaging and scRNA-seq along with ST data could achieve a higher cellular resolution. (15). Robust Cell Type Decomposition (RCTD), a computational method that attains cell type profiles from scRNA-seq to decompose low-resolution cell mixture spots into discrete cells. (16) In addition, several *in silico* deep learning-based computational algorithms have also been introduced, without any scRNA-seq dataset or preselected genes. SPICEMIX based on probabilistic and latent variable modeling for analysis of ST dataset can refine cell state with revealing spatially variable features. (17) BayesSpace, a Bayesian statistical method that uses the information from spatial neighborhoods could enhance the resolution of the ST dataset. (18)

Herein, we devised a novel methodology, that integrates the I-B and B-B ST dataset, could extend the gene coverage in the I-B ST dataset while maintaining the high resolution: GeneDART, an application of deep learning to extent gene coverage of I-B ST. Notably, GeneDART used the B-B ST dataset to complement the gene coverage range in I-B ST, rather than the scRNA-seq dataset. In utilizing the scRNA-seq dataset for the previously reported methods for ST dataset integration, scRNA-seq imputation methods have been used because of ‘dropout’ events during the technical procedures and low read counts of each cell in scRNA-seq data, (19) aiming at imputing ‘false-negative’ detection of scRNA-seq data. Meanwhile, our GeneDART predicts the genes which are not included in the I-B ST dataset. In this aspect, the purpose of the model is largely distinctive from currently reported methods. Furthermore, GeneDART is noteworthy in its elements that make up the methodology. The domain adaptation based on adversarial loss to integrate different feature domain data. An adversarial loss was used for the domain adaptation to integrate two different feature domains: B-B and I-B data. Also, it is possible that GeneDART could provide an internal quality check process using a random gene set whether this model is fit and functions properly for predicting target features. Here, we monitored the performance based on the ‘*Genes for Test*’ from the common gene set, playing a role as a validation strategy of GeneDART,

Though, some issues need to be further addressed: we confirmed that this method effectively extended gene coverage in I-B ST, specifically a newly introduced technology, Xenium. It should be needed to be validated that this methodology could be applied to another I-B ST platform, such as CosMX (NanoString, WA, USA). Because these methods use different principles to measure gene expression, different data distributions can affect the performance and accordingly another tuning process of GeneDART may be needed. Nevertheless, this current approach is meaningful in that it provides a milestone leveraging advantages of B-B and I-B-based ST methods through domain adaptation. Secondly, it may be crucial to select a proper B-B ST dataset obtained from the same or very similar tissues and/or organs when training a model in applying GeneDART, because an erroneous gene expression would be predicted when B-B ST data are incompatible with I-B ST data.

Herein, we presented a deep learning model with adversarial loss of ST data by conjoining both technologies, thereby extending gene coverage in I-B ST with a highly preserved spatial resolution in a complementary role.

## Acknowledgments

We thank all members of Portrai, Inc. for the discussion and technical support.

## References

1. Rao, A., Barkley, D., França, G.S. and Yanai, I. (2021) Exploring tissue architecture using spatial transcriptomics. Nature, 596, 211–220.

2. Moses, L. and Pachter, L. (2022) Museum of spatial transcriptomics. Nat. Methods, 19, 534–546.

3. Williams, C.G., Lee, H.J., Asatsuma, T., Vento-Tormo, R. and Haque, A. (2022) An introduction to spatial transcriptomics for biomedical research. Genome Med., 14, 68.

4. Lewis, S.M., Asselin-Labat, M.L., Nguyen, Q., Berthelet, J., Tan, X., Wimmer, V.C., Merino, D., Rogers, K.L. and Naik, S.H. (2021) Spatial omics and multiplexed imaging to explore cancer biology. Nat. Methods, 18, 997–1012.

5. Holt, C.E. and Bullock, S.L. (2009) Subcellular mRNA localization in animal cells and why it matters. Science, 326, 1212–1216.

6. Borm, L.E., Albiach, A.M., Mannens, C.C.A., Janusauskas, J., Özgün, C., Fernández-Garc ía, D., Hodge, R., Lein, E.S., Codeluppi, S. and Linnarsson, S. (2022) Scalable in situ single-cell profiling by electrophoretic capture of mRNA. bioRxiv, 2022.2001.2012.476082.

7. Codeluppi, S., Borm, L.E., Zeisel, A., La Manno, G., van Lunteren, J.A., Svensson, C.I. and Linnarsson, S. (2018) Spatial organization of the somatosensory cortex revealed by osmFISH. Nat. Methods, 15, 932–935.

8. Wolf, F.A., Angerer, P. and Theis, F.J. (2018) SCANPY: large-scale single-cell gene expression data analysis. Genome Biol., 19, 15.

9. Hildebrandt, F., Andersson, A., Saarenpää, S., Larsson, L., Van Hul, N., Kanatani, S., Masek, J., Ellis, E., Barragan, A., Mollbrink, A. et al. (2021) Spatial Transcriptomics to define transcriptional patterns of zonation and structural components in the mouse liver. Nat. Commun., 12, 7046.

10. Andersson, A., Larsson, L., Stenbeck, L., Salmén, F., Ehinger, A., Wu, S.Z., Al-Eryani, G., Roden, D., Swarbrick, A., Borg, Å. et al. (2021) Spatial deconvolution of HER2-positive breast cancer delineates tumor-associated cell type interactions. Nat. Commun., 12, 6012.

11. Hunter, M.V., Moncada, R., Weiss, J.M., Yanai, I. and White, R.M. (2021) Spatially resolved transcriptomics reveals the architecture of the tumor-microenvironment interface. Nat. Commun., 12, 6278.

12. Heindl, A., Nawaz, S. and Yuan, Y. (2015) Mapping spatial heterogeneity in the tumor microenvironment: a new era for digital pathology. Lab. Invest., 95, 377–384.

13. Lohoff, T., Ghazanfar, S., Missarova, A., Koulena, N., Pierson, N., Griffiths, J.A., Bardot, E.S., Eng, C.L., Tyser, R.C.V., Argelaguet, R. et al. (2022) Integration of spatial and single-cell transcriptomic data elucidates mouse organogenesis. Nat. Biotechnol., 40, 74–85.

14. Rodriques, S.G., Stickels, R.R., Goeva, A., Martin, C.A., Murray, E., Vanderburg, C.R., Welch, J., Chen, L.M., Chen, F. and Macosko, E.Z. (2019) Slide-seq: A scalable technology for measuring genome-wide expression at high spatial resolution. Science, 363, 1463–1467.

15. Ji, A.L., Rubin, A.J., Thrane, K., Jiang, S., Reynolds, D.L., Meyers, R.M., Guo, M.G., George, B.M., Mollbrink, A., Bergenstr åhle, J. et al. (2020) Multimodal Analysis of Composition and Spatial Architecture in Human Squamous Cell Carcinoma. Cell, 182, 497-514.e422.

16. Cable, D.M., Murray, E., Zou, L.S., Goeva, A., Macosko, E.Z., Chen, F. and Irizarry, R.A. (2022) Robust decomposition of cell type mixtures in spatial transcriptomics. Nat. Biotechnol., 40, 517–526.

17. Chidester, B., Zhou, T., Alam, S. and Ma, J. (2023) SpiceMix enables integrative single-cell spatial modeling of cell identity. Nat. Genet., 55, 78–88.

18. Zhao, E., Stone, M.R., Ren, X., Guenthoer, J., Smythe, K.S., Pulliam, T., Williams, S.R., Uytingco, C.R., Taylor, S.E.B., Nghiem, P. et al. (2021) Spatial transcriptomics at subspot resolution with BayesSpace. Nat. Biotechnol., 39, 1375–1384.

19. Dai, C., Jiang, Y., Yin, C., Su, R., Zeng, X., Zou, Q., Nakai, K. and Wei, L. (2022) scIMC: a platform for benchmarking comparison and visualization analysis of scRNA-seq data imputation methods. Nucleic Acids Res., 50, 4877–4899.

